# A neurophysiological study of noun-adjective agreement in Arabic: The impact of animacy and diglossia on the dynamics of language processing

**DOI:** 10.1101/729855

**Authors:** Ali Idrissi, Eiman Mustafawi, Tariq Khwaileh, R. Muralikrishnan

**Affiliations:** Qatar University; Max Planck Institute for Empirical Aesthetics

**Keywords:** Arabic, agreement, animacy, diglossia, ERPs, N400/P600

## Abstract

We used event-related brain potentials to identify the neurophysiological responses of Arabic speakers to processing full and deflected agreement in plural noun-adjective constructions in Standard Arabic. Under full agreement, an adjective fully agrees in number and gender with a preceding plural noun, but only when this noun is human, while it is systematically marked feminine singular when the noun is non-human under deflected agreement. We recorded grammaticality judgment and ERP responses from 32 speakers of Arabic to sentences violating full and deflected agreement and their well-formed counterparts. The participants were relatively fast and accurate in judging *all* the sentences, although violations, especially deflected agreement violations, were not always deemed ungrammatical. However, the ERP responses show a differential processing of human versus non-human violations. Violations of full agreement involving human nouns elicited larger N400 and P600 components than violations of deflected agreement involving non-human nouns, whose ERP signatures were hardly distinguishable from those of their acceptable counterparts. Our results present evidence for animacy (more specifically, humanness) and inter-dialect effects on language processing. We argue that violations of Standard Arabic deflected agreement are not treated as outright violations because non-human referents permit both full and deflected agreement in Spoken Arabic. We discuss these results in light of the ERP literature on agreement processing and the role of animacy/humanness in grammar, and highlight the potential effect of diglossia on the architecture of the mental grammar of Arabic speakers.

## 1 Grammatical agreement

Structural dependencies between different constituents in a sentence are typically expressed by means of grammatical agreement. In many languages, a semantic and/or syntactic property of a constituent is overtly marked on one (or more) other constituent in the same sentence (Corbett, 2006; Nichols & Bickel, 2013; Steele, 1978; Wunderlich, 2015). This systematic covariance between a semantic or syntactic property of a word and the formal features of another is the hallmark of grammatical agreement. A common example is subject-verb agreement, illustrated in (1) with sentences from Standard Arabic.^1^ The processing of subject-verb agreement has been extensively studied in many languages (Angrilli et al., 2002; Hinojosa, Martín-Loeches, Casado, Muñoz, & Rubia, 2003; Münte, Szentkuti, Wieringa, Matzke, & Johannes, 1997; Nevins, Dillon, Malhotra, & Phillips, 2007; Osterhout & Mobley, 1995; Palolahti, Leino, Jokela, Kopra, & Paavilainen, 2005).

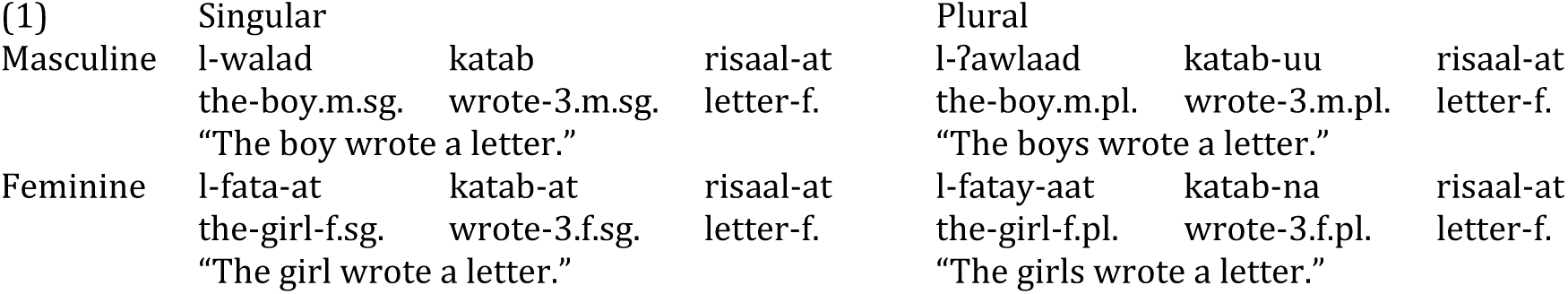

Agreement relations can obtain within the same clause, such as between a subject and a verb, illustrated in (1), or between a noun and a predicate adjective, as shown in (2).

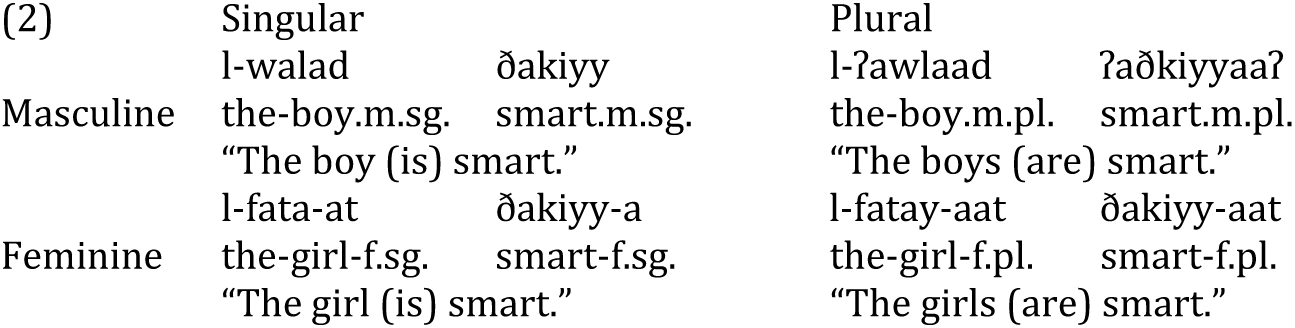

Agreement relations can also obtain within a noun phrase, namely between a noun and a determiner (3a-d) or a noun and a modifying adjective (3e-h).

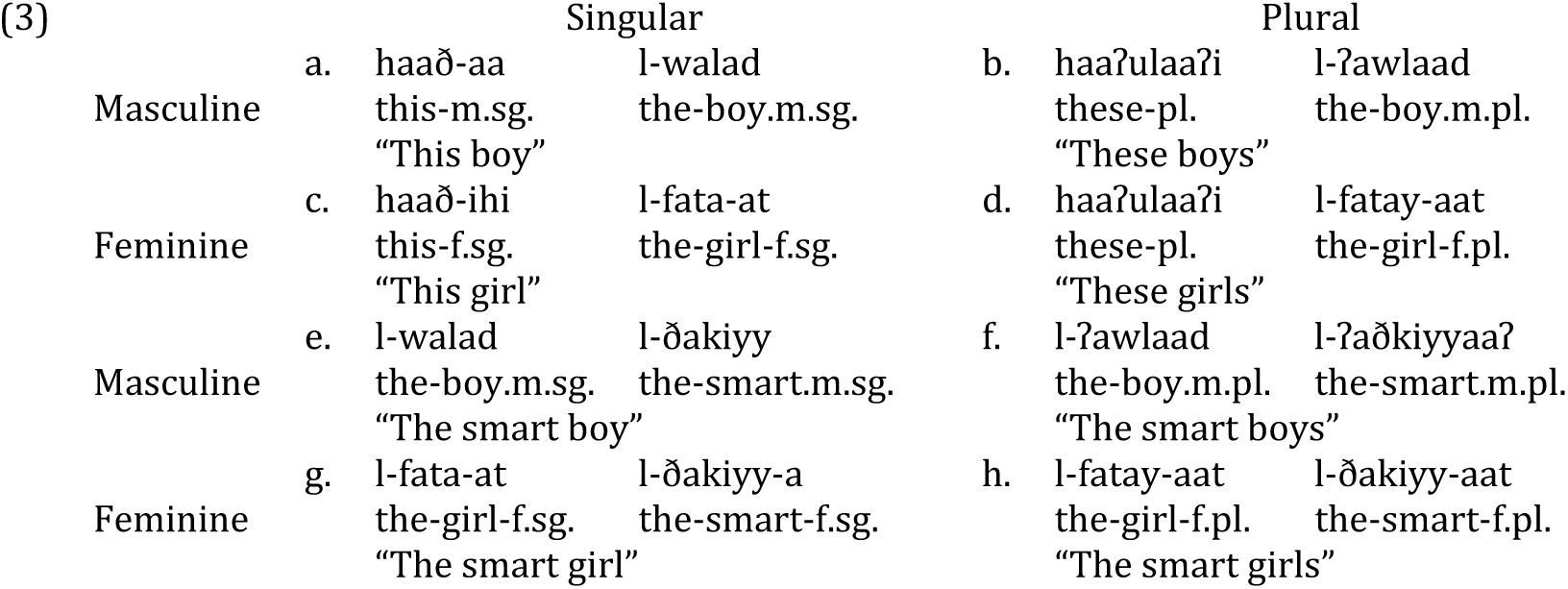

The neural correlates of the processing of noun-adjective agreement have been studied, although to a lesser extent than subject-verb agreement (see Kutas & Hillyard (1983) for English, and Barber & Carreiras (2005) and Martin-Loeches, Nigbur, Casado, Hohlfeld, & Sommer (2006), for Spanish) and noun-determiner agreement (see Hagoort (2003) for Dutch, and Barber & Carreiras (2005) for Spanish).

Because it is obvious on the controller (i.e., the trigger of agreement), the information repeated on structurally related words in the same sentence can be considered redundant. Yet, this redundancy is not only present cross-linguistically, but languages vary in terms of the number of features involved and the number of targets affected. For example, while both number and gender are covaried in Arabic (as shown in (1) above), in English only number is (as shown in (4)).

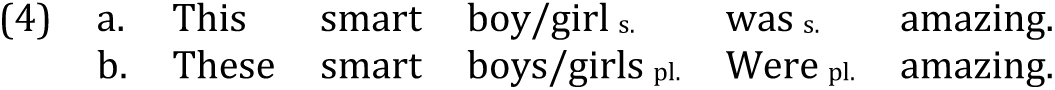

Additionally, while agreement is expressed on only two of the five words in the English sentences in (4), the demonstrative and the verb, it is overt on four out of the five words in the Arabic sentences in (5): the demonstrative, the two adjectives, and the verb.

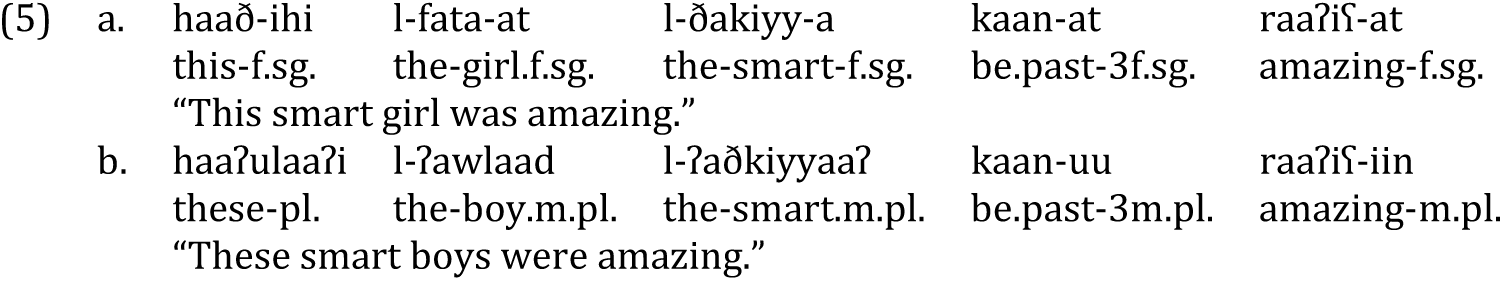

Given that structural dependencies are generally morphosyntactically marked, and given their importance for the construction of sentence meaning, processing grammatical agreement must be critical in language comprehension. It should be even more so in languages like Arabic with relatively free word order and rich inflectional morphology (Deutsch & Bentin, 2001; Molinaro, Barber, & Carreiras, 2011). During the computation of sentence meaning in Arabic, the processor must thus verify that structurally dependent words in a given sentence bear the right inflectional markers.

In this study we are concerned with how number and gender features are processed in Standard Arabic noun-adjective agreement. Noun-adjective agreement in Standard Arabic is characterized by the presence of the canonical pattern of agreement (which we call ‘full’ agreement) along with an unusual pattern of grammatical *dis*agreement (commonly called ‘deflected’ agreement) (Ferguson, 1989). Our goal is three-fold: (i) determine the general neurophysiological patterns associated with noun-adjective agreement in Arabic, (ii) examine the role animacy (specifically, humanness) plays in Arabic noun-adjective agreement processing, and, given that our participants are all diglossic speakers, (ii) explore the possible impact of interference from Spoken Arabic during Standard Arabic sentence processing.

The paper is organized as follows. Section 2 introduces the phenomena of full agreement and deflected agreement in Standard Arabic noun-adjective constructions. Section 3 provides a brief review of the ERP literature on the processing of grammatical agreement. The questions and hypotheses underlying the current study are given in Section 4. The methodology and the results are given in sections 5 and 6 respectively. The results are discussed in Section 7, and Section 8 concludes the paper.

### 2 Noun-adjective agreement patterns in Arabic

Arabic shows both the typical pattern of canonical agreement, where features of the trigger and target match, and an unusual pattern of disagreement where the target morphologically disagrees with the *masculine plural* trigger and always appears in the *feminine singular*. The distinction between these two patterns arises only when the controller is plural and are referred to as full or strict agreement (henceforth FA) and deflected agreement (henceforth DA), respectively (Ferguson, 1989). Along with agreement asymmetry observed in the VSO order in Standard Arabic (Aoun, Benmamoun, & Choueiri, 2010; Fassi-Fehri, 1981), DA constitutes an empirically and theoretically intriguing phenomenon in Arabic, as evidenced by the amount of descriptive and theoretical work that has been devoted to it (Belnap, 1993; Belnap & Haeri, 1997; Bettega, 2017; Dali & Mathieu, 2016; D’Anna, 2017; Ferguson, 1989; Procházka & Gabsi, 2017; Ritt-Benmimoun, 2017).

Importantly, the distribution of the FA and DA patterns is governed by animacy, and more specifically, by humanness. In fact, when the trigger noun is nonhuman (a creature (6c) or an inanimate entity (6d)), it systematically imposes feminine singular agreement, i.e. DA, on the target, even when it itself is masculine. By contrast, when the trigger is human, as in(6a) and (6b), FA applies. In (6a-b) inflectional morphology is co-varied as expected under FA, but it is varied in such a way that the feature/s of the target is/are in *complete* mismatch with the feature/s of the non-human trigger in (6c-d).

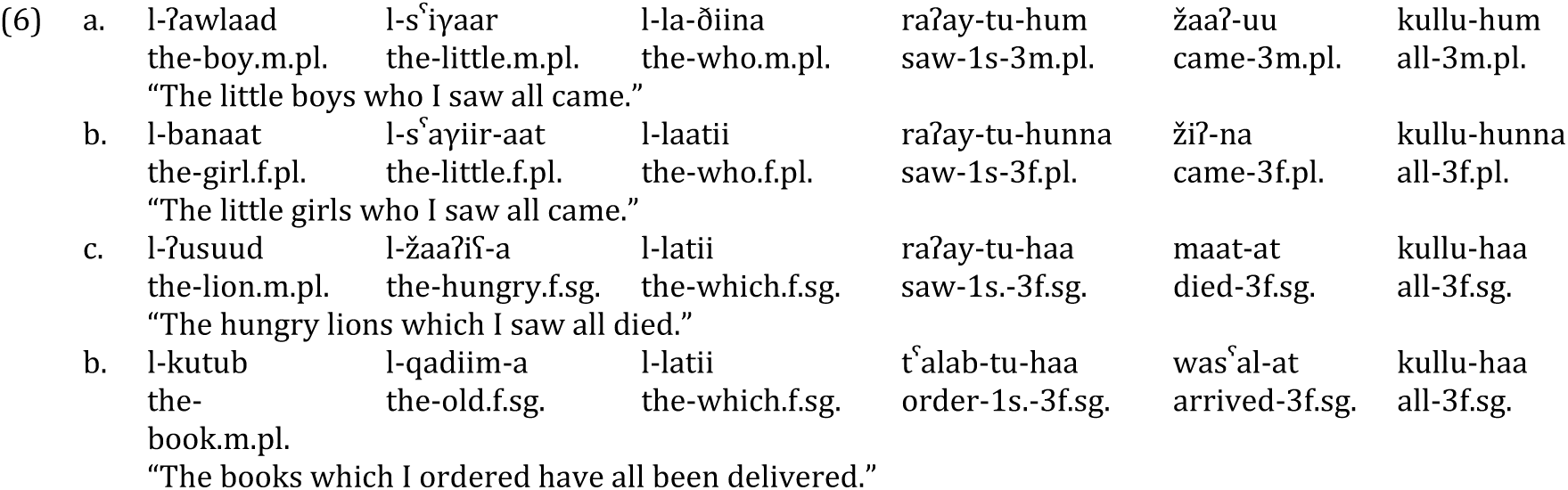

Interestingly, while Standard Arabic strictly adheres to this distribution of FA and DA, modern spoken varieties of Arabic allow both FA and DA where only DA is expected in Standard Arabic, especially with non-human referents, and to greatly lesser extent, human broken plural referents (Belnap, 1993; Bettega, 2017; Ferguson, 1989). Thus, both FA and DA are grammatical in Spoken Arabic although there may be a preference for one or the other. In some cases one pattern is indeed more unmarked/default for one construction than for the other, and in others, the two patterns can be in free variation ((Ambros, 1977). The difference between Standard Arabic and Spoken Arabic plural noun agreement patterns is summarized in (7).

|  |  |  |  |
| --- | --- | --- | --- |
| (7) |  | Controller noun | Agreement pattern |
|  | Standard Arabic | Human | Full agreement |
|  |  | Non-human | Deflected agreement |
|  | Spoken Arabic | Human | Full agreement |
|  |  | Nonhuman | Deflected agreement / Full agreement |

Belnap (1993) observes that in Cairene Arabic, human sound plurals never take DA, human broken plurals only rarely do so (16%), and inanimate sound and broken plurals predominantly take DA (91% and 92% respectively). As for non-human animate plurals (the class of plurals we studied in this experiment), they show a 33% incidence of FA. This author also observes that the distribution of DA and FA with non-human nouns is sensitive to the distance/proximity between the trigger and the target. Specifically, DA becomes less used (and is replaced with FA) as the distance between the locus of agreement and the target becomes larger. More or less the same distribution has been reported in Omani Arabic (Bettega, 2017), Tunisian Arabic (Procházka & Gabsi, 2017), and Libyan Arabic (D’Anna, 2017).

In this study we investigate how FA and DA are computed in Standard Arabic noun-adjective combinations of the type illustrated in (6). The interest of this pattern lies in that it provides an ideal ground to see how the processor resolves the conflict between surface feature mismatch (syntactic agreement) and the underlying feature match driven by semantics (semantic agreement).^2^ This pattern also allows us to test the interaction of animacy with morphosyntax, and the time course of their computation. We may also be able to explore how distinct the cognitive mechanisms underlying the processing of grammatical properties, such as number and gender, are from those associated with the computation of conceptual/semantic features, such as animacy. Standard Arabic offers a case where the way grammar morphologically encodes these features does not coincide with how the world is categorized, with cases of a clear mismatch between morphosyntactic features, on the one hand, and plurality and biological gender, on the other. Finally, we can indirectly provide some insight into the impact of diglossia on language processing and the overall architecture of the mental grammar of Arabic speakers, since most Spoken Arabic varieties display a different and more flexible agreement pattern than Standard Arabic, with FA being often used in alternation with DA with non-human referents in some varieties and almost instead of it in others.

The ERP literature on the processing of agreement is abundant. However, most of it deals with clausal subject-verb agreement, with agreement within DP (i.e., concord) being less studied. Additionally, a look at Molinaro et al.’s (2011) review shows that the majority of languages where the processing of agreement has been addressed are all Indo-European, except for two studies on Finnish (see Leinonen, Brattico, Järvenpää, & Krause, 2008; Palolahti et al., 2005), and one on Hebrew (see Deutsch & Bentin, 2001). There is therefore need for typologically different languages to be investigated, not only for the sake of contributing to the growing body of research in the field of agreement processing, but also for the sake of cross-linguistic validation of available results (Díaz, Sebastián-Gallés, Erdocia, Mueller, & Laka, 2011). Moreover, we have an opportunity to shed light on an idiosyncratic feature in Arabic grammar which presents an intriguing agreement pattern, whose structure is governed by animacy and whose processing may be sensitive to the nature of the diglossic competence of contemporary Arabic speakers.

## 3 ERPs of agreement processing

### 3.1 Main language-related ERP components

ERPs have proven an excellent technique to investigate the processing of grammatical agreement during language comprehension (Barber, Salillas, & Carreiras, 2004; Hagoort, 2003; Hagoort, Brown, & Groothusen, 1993; Kutas & Hillyard, 1983). Most ERP agreement experiments use the violation paradigm which consists of manipulations of agreement comparing constructions abiding by agreement rules with their ungrammatical counterparts where agreement is (partially or fully) violated. Mean amplitude differences between the different conditions at different scalp sites, and at different latencies from the onset of the critical word are then taken to reveal the mechanisms involved in agreement processing. Three major ERP components have emerged in the literature as being associated with the processing of agreement (Friederici, 1995).

The first among these is the N400, which is a negative-going neurophysiological component which normally peaks between 300 and 450 ms after the onset of the stimulus and shows a centroparietal topographical distribution. It increases in amplitude as a function of the amount of cognitive resources deployed during single word processing, and it is also reported to index processes related to sentence-level expectancy (Alday, Schlesewsky, & Bornkessel-Schlesewsky, 2017; Lau, Phillips, & Poeppel, 2008; Molinaro et al., 2011). In a broader theory of the neurobiology of negativity, the N400 is also argued to reflect prediction errors (Bornkessel-Schlesewsky & Schlesewsky, 2019).

The second, LAN (Left Anterior Negativity), is also a negative going component which overlaps with N400 temporally but shows a rather left-frontal scalp distribution. It has been taken to index morphosyntactic processing over ungrammatical constructions (Osterhout & Mobley, 1995). A distinction has been made between an early LAN (at about 100 ms) associated with early morphosyntactic violations, and a later LAN linked to later morphosyntactic processing (Friederici, 2004). Because LAN has not been consistently reported, its etiology and exact status remain less well-established (Tanner & van Hell, 2014).

The third component, which has been consistently found in studies involving agreement violations, is the P600. P600 is a positive going wave which peaks between 500 and 1000 ms and shows a larger distribution over posterior scalp electrodes. It is said to correlate with late syntactic analysis and reanalysis/repair of syntactic anomalies (Bornkessel-Schlesewsky & Schlesewsky, 2008; Osterhout & Mobley, 1995) and more generally with conflict-monitoring processes (van de Meerendonk, Kolk, Chwilla, & Vissers, 2009). A distinction has been made between two types of P600 (Molinaro et al., 2011). First, there is the early P600, which peaks between 500 and 750 ms and shows a rather broad scalp distribution and correlates with the difficulty to integrate a given word within its syntactic context. Then, the later P600 is a component that peaks between 750 and 1000 ms and is restricted mainly to posterior electrodes and is argued to reflect reanalysis or repair routines.

### 3.2 ERPs of number and gender agreement

ERP studies of subject-verb agreement report a biphasic LAN – P660 pattern for number mismatches (Mancini, Molinaro, Rizzi, & Carreiras, 2011; Molinaro, Vespignani, & Job, 2008). Likewise, determiner-noun and noun-adjective agreement mismatches have been reported to yield a LAN – P600 pattern; but Barber & Carreiras (2003, 2005) reported an N400 effect in response to both number and gender agreement violations involving adjectives and nouns, and a LAN and a P300 for gender violations only.

Gender agreement violations within the clause have been shown to trigger a LAN – P600 pattern (Molinaro et al., 2011, 2008), and a study on gender agreement in Spanish by O’Rourke & Van Petten (2011) reported a LAN response to gender agreement violations regardless of the distance between the disagreeing elements. However, some studies did not report a LAN at all (Hagoort, 2003; Popov & Bastiaanse, 2018; Wicha, Moreno, & Kutas, 2004). In these studies, and article-noun and noun-adjective number and gender agreement elicited a P600 but no LAN. A recent study by Mancini et al. (2011) reported a different distribution of the late-positivity in addition to a qualitative difference in the negativities, namely a LAN effect for number and an N400 effect for person.

Using eye-tracking and ERPs, Deutsch & Bentin (2001) manipulated subject–predicate gender agreement in Hebrew. There was no overt gender marker on the noun, gender being signaled with word internal structure. The authors reported longer reading times for disagreeing than for agreeing predicates when gender was overtly marked on the adjective. They also showed that gender violations led to larger amplitude of eLAN, a later N400, and a P600.

In a recent study on Arabic, Melebari (2017) studied FA and DA in determiner – noun constructions in Standard Arabic. Her behavioral results show that participants were highly accurate in the singular conditions, and relatively so in the plural conditions. It indeed emerges that the plural mismatch conditions were relatively less easy to decide on. Melebari’s ERP results show that in the singular, violations elicited negative-going ERP responses for both human and non-human conditions. In the plural, a polarity reversal for non-human nouns was observed: plural human violations (like their singular counterparts) elicited a negative going violation response; whereas plural nonhuman violations elicited a relative positivity. In the 900-1100 ms and 1100-1300 ms time-windows, non-human plurals elicited a mirror image pattern to the other correct/violation comparisons.

In sum, agreement violations have generally been said to elicit a LAN (in the 300 - 500 ms time window) followed by a late positivity (P600) (in the 500 - 700 ms window and often later). However, a few studies reported an N400, instead of a LAN, followed by a P600 (Díaz et al., 2011; Kaan, Harris, Gibson, & Holcomb, 2000; Zawiszewski, Santesteban, & Laka, 2016).

## 4 The current study

From the brief review in the preceding section, there seems to be a somewhat clear picture of the neural correlates of morphosyntactic incompatibility between the controller and target in various agreement configurations and domains. The goal of this study is to explore a unique case where this incompatibility consists of a mismatch in one case (violation of FA but adherence to DA) and an ungrammatical match in the other (violation of DA but adherence to FA). This is done in an equally unique context, diglossia, in which the linguistic system of the participants’ spoken variety allows alternative agreement patterns, a situation which should plausibly lead to some form of processing flexibility where the parser has “loose” patterns/rules at its disposal. As a consequence of this flexibility, what is potentially ungrammatical in one system may be tolerated (to some degree) during sentence processing because it is grammatical in the other.

We thus address two major questions in this paper. First, how are FA and DA processed in Arabic, assuming that they are different surface manifestations of the same general principle/rule: Agreement Rule? Specifically, are they treated the same, or do they call for different processing routines? Second, what is the effect of inter-dialectal (or more accurately, inter-variety) interference on DA processing given that it alternates with FA in Spoken Arabic?

Research on agreement has so far focused on a small set of languages and on typical cases of agreement violations. Except for work by Mancini and colleagues in Spanish (Mancini, Molinaro, Rizzi, & Carreiras, 2011) and Basque (Mancini, Massol, Duñabeitia, Carreiras, & Molinaro, 2019), there have been no studies looking into idiosyncratic cases of agreement. (Mancini et al., 2011) studied a relatively similar phenomenon in Spanish, which they call “unagreement”. However, the mismatch phenomenon they studied involves person only and concerns clausal subject-verb dependency; and their results show that person unagreement triggered a left posterior negativity followed by a more central negativity, with no P600 effect. Therefore, our study not only contributes to the cross-linguistic scope of ERP studies by focusing on a highly understudied language, Arabic; but it also constitutes the first study to systematically examine an unusual gender and number agreement pattern, which is actually a form of syntactic disagreement.

We hypothesize that FA violations should not be tolerated due to the relative importance of humanness in the agreement system of both Standard Arabic and Spoken Arabic varieties. The parser should detect the gender and number agreement mismatch between the human noun and the adjective, and a negativity effect (most likely N400) should ensue. We also expect that DA violations be amnestied by the parser due to variation in the agreement system of Spoken Arabic in which FA is a common option used along with DA with non-human controllers. So, a smaller negativity or no negativity at all should be observed for DA violations, compared to FA violations. Finally, for FA violations, although not tolerated, an attempt to tolerate them should be observed due to the prevalence of non-human plurals, and therefore of DA in the language. It follows that when faced with a FA violation, the parser should attempt a DA reading leading to a concomitant delay in the activation of repair and/or conflict-monitoring processes briefly mentioned earlier (van de Meerendonk et al., 2009). It remains to be seen whether this will be reflected in the P600 latency.

## 5 Experimental procedures

### 5.1 Participants

Thirty-two Qatar University female students participated in the experiment. All participants were right-handed native Arabic speakers, with normal or corrected-to-normal vision and normal hearing. None of them reported history of brain injury or brain surgery, or any form of linguistic or cognitive impairment. All participants gave their written informed consent prior to their participation and were compensated for their time. The experiment, participant recruitment and compensation were approved by Qatar University Institutional Review Board. Six participants had to be excluded from the analyses because of excessive EEG artefacts and/or too many errors in the behavioral control task, leaving a total of 26 participants for the analysis.

### 5.2 Materials

All critical sentences were of the form *noun-adjective-verb–prepositional phrase*. The subject nouns were either human or animate non-human (animal) masculine plural; the adjective was either in correct agreement (FA or DA depending on whether the noun was human or nonhuman) or it violated agreement. The verbs were all transitive and in the past tense. As a first step, 36 human nouns and 36 non-human animate nouns were used with an appropriate adjective to construct acceptable sentences. The violation conditions were generated from each of the thus resulting 72 acceptable sentences such that the agreement marking on the adjective was in violation of Standard Arabic agreement rules, showing feminine singular for human nouns (violation of FA) and masculine plural for non-human nouns (violation of DA). This resulted in a total of 36 sets of sentences in four critical conditions, with 144 critical sentences. The four conditions are exemplified below (H = Human, N = Nonhuman, A = Acceptable (or Grammatical), and V = Violation). Ungrammaticality is indicated with an asterisk. The Arabic script is read from right to left, and the phonetic transcriptions are read from left to right.

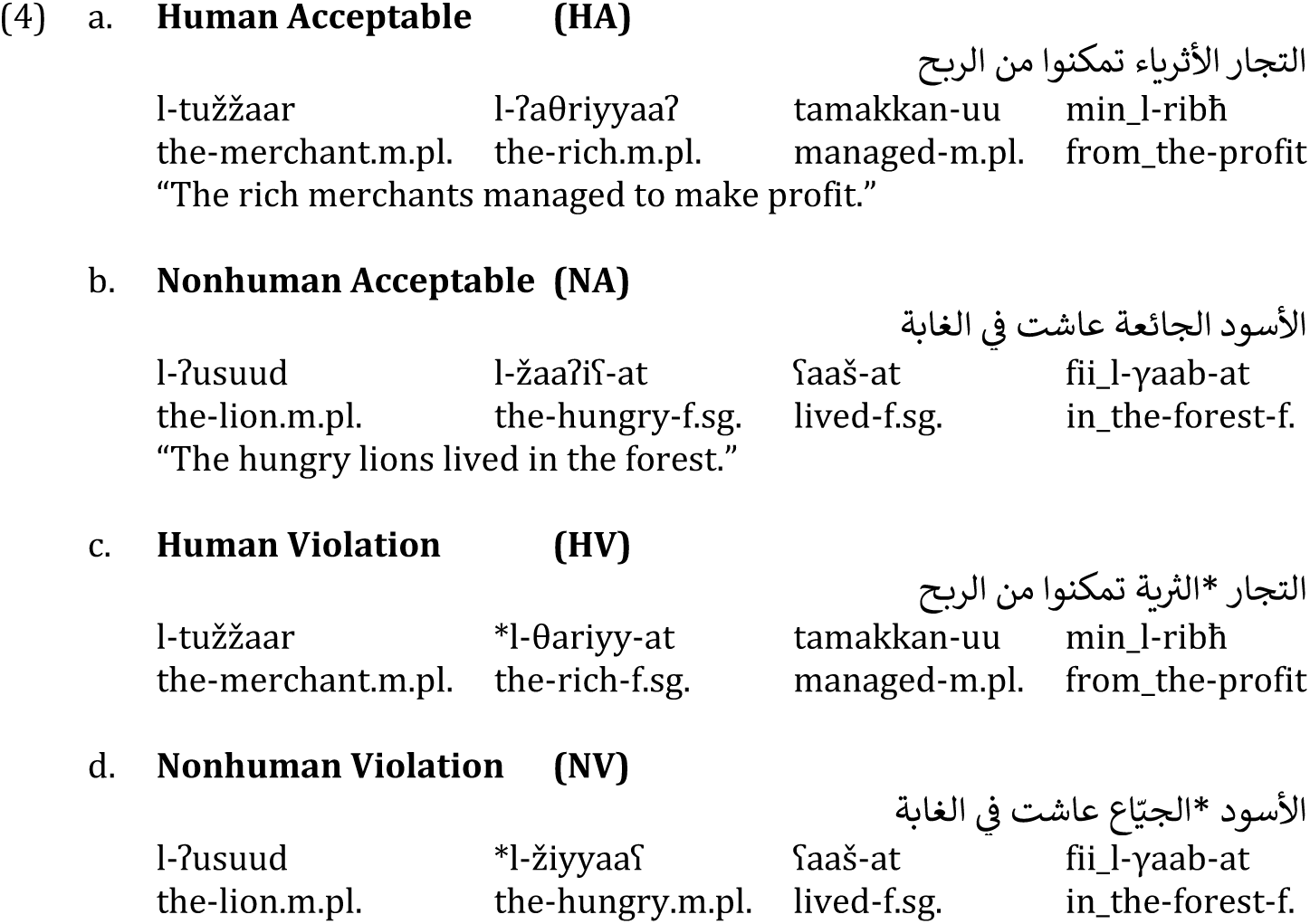

HA sentences adhere to the canonical agreement (i.e. FA), and NA sentences to grammatical disagreement (i.e. DA); whereas HV and NV violate FA and DA, respectively.

Fillers from an unrelated design involving transitive and subject-dropped structures and a few semantic anomalies were distributed into two unique sets and interspersed with the critical stimuli such that the sentence structure was not predictable. The overall number of grammatical versus ungrammatical sentences was counterbalanced in the resulting stimuli sets consisting of 144 targets and 288 fillers. The two sets were each pseudo-randomized twice to obtain four stimulus lists, one of which was used for every participant. The presentation of the randomized lists was counterbalanced across participants. Where possible in the critical sentences, we included both broken and sound plurals as critical nouns; and when the adjective was plural, we used 18 broken plural adjectives and 18 sound plural ones. When the adjective was feminine singular, the only form possible was feminine by suffixation. The length of the critical items (the trigger and the target) was balanced across all 4 conditions.

### 5.3 Tasks

Given the use of a violation paradigm, an acceptability judgement task followed the presentation of each stimulus sentence, which required a ‘yes’ or ‘no’ answer. In addition, in order to ensure that participants read the sentences attentively, a probe-word detection task followed the acceptability judgement task. The probe task was constructed in such a way that an equal number of trials required ‘yes’ or ‘no’ as answers. If the probe word appeared in the preceding stimulus sentence, this required a ‘yes’ answer, whereas if it did not, the required answer was a ‘no’. Crucially, the word position from which the probe word was chosen was equiprobable across the experiment as well as within each condition, which meant that participants had to be very attentive throughout the stimulus sentence presentation in order to perform the task correctly.

### 5.4 Procedure

The experiment was conducted at the Neurocognition of Language Lab at Qatar University. Participants filled an Edinburgh-Handedness questionnaire in Arabic, and onlydominant right-handers were accepted for participation. They received printed instructions about the experiment and the tasks they had to perform. Stimuli were presented using the Presentation software (www.neurobs.com) that recorded, among other things, the trial number, reaction time and the button responses. The participants were instructed to read as normally and carefully as possible. The brightness and contrast settings of the monitor were maintained the same for all the participants.

Once the electrode cap was set up, participants were seated on a comfortable chair in a sound-proof chamber and requested to avoid abrupt and drastic movements, especially movements of the head. Then, the so-called resting EEG was recorded for possible frequency-based EEG analyses later, where the participant had to sit still for two minutes with no specific task to perform. Two more minutes of resting EEG was recorded, but this time the participants had to close their eyes. After a short pause, the experimental session began, which consisted of a short practice followed by the actual experiment.

The structure of each trial in the experiment was as follows. The flat-screen LCD monitor was clear before the trial started. A fixation asterisk was shown in the center of the screen for 500 ms, after which the screen became blank for 100 ms. Then the rapid serial visual presentation of the stimulus sentence started. Each word appeared in the center of the screen and remained for 600 ms, after which the screen became blank for 100 ms before the next word appeared. When they consisted of two orthographically separate words, prepositional phrases were presented for 750 ms. After the last word of the stimulus sentence was presented, the screen was blank for 500 ms. Following this, a pair of smileys appeared on the screen, prompting the participant to judge the acceptability of the sentence. After a maximum of 2000 ms or after a button press, whichever was earlier, the screen became blank again for 500 ms. A time-out was registered when no button was pressed within 2000 ms. A probe word then appeared in the middle of the screen for a maximum of 2000 ms, within which the participant had to detect whether the word appeared in the preceding stimulus sentence or not. When no button was pressed within 2000 ms, a time-out was registered. At the end of the trial, the screen became blank for a 1500 ms inter-stimulus interval before the next trial started.

Before the actual experiment began, there was a short practice session consisting of twelve trials, which was aimed at helping participants to get used to the task and feel comfortable with the pace of the trials and the blinking regime. For any given participant, none of the experimental stimuli occurred in their practice phase. The task in the practice session was identical to that of the experiment phase. The EEG of the participants was not recorded in the practice phase. In the main phase of the experiment, one of the four pseudo-randomized sets of materials mentioned above was chosen to be presented in 12 blocks of 36 trials each. There was an equal number of probe words that required a ‘Yes’ or ‘No’ answer in each block. For the sake of counterbalancing for any right-dominance effects, half of the participants had the ‘Yes’ button on the right side, and the other half had it on the left side. The ‘Yes’ button being on the right or left was also counterbalanced across the stimuli sets. There was a short pause between blocks. Resting EEG was again recorded at the end of the experimental session.

### 5.5 EEG recording, pre-processing and statistical analysis

The EEG was recorded by means of 25 AgAgCl active electrodes fixed at the scalp by means of an elastic cap (Easycap GmbH, Herrsching, Germany). AFZ served as the ground electrode. Recordings were referenced to the left mastoid, but re-referenced to the average of linked mastoids offline. The electrooculogram (EOG) was monitored by means of electrodes placed at the outer canthus of each eye for the horizontal EOG and above and below the participant’s right eye for the vertical EOG. Electrode impedances were kept below appropriate levels to ensure a good quality signal with minimal noise. All EEG and EOG channels were amplified using a BrainAmp amplifier (Brain Products GmbH, Gilching, Germany) and recorded with a digitization rate of 250 Hz. The EEG data thus collected was pre-processed for further analysis using a 0.3−20Hz band pass filter in order to remove slow signal drifts. The statistical analyses were performed on this data, but an 8.5 Hz low-pass filter was further applied on the data for achieving smoother ERP plots.

ERPs were calculated for each participant from 200 ms before the onset of the verb until 1200 ms after onset (so −200 ms to 1200 ms). These were averaged across items per condition per participant before computing the grand-average ERPs across participants per condition. Repeated-measures analyses of variance (ANOVAs) were computed for the statistical analysis of the ERP data, involving the within-participants factors Humanness (HN) and Agreement (AT) for mean amplitude values per time-window per condition in 4 lateral regions of interest (ROIs) and 6 midline ROIs. The lateral ROIs were defined as follows: LA, comprised of the left-anterior electrodes F7, F3, FC5 and FC1; LP, comprised of the left-posterior electrodes P7, P3, CP5 and CP1; RA, comprised of the right-anterior electrodes F8, F4, FC6 and FC2; and RP, comprised of the right-posterior electrodes P8, P4, CP6 and CP2. Each of the 6 midline electrodes FZ, FCZ, CZ, CPZ, PZ and POZ constituted an individual midline ROI of the same name respectively.

The statistical analysis of the ERP data was carried out in a hierarchical manner, that is to say, only interactions that were at least marginally significant were resolved. To avoid excessive type 1 errors due to violations of sphericity, the correction of Huynh & Feldt (1970) was applied when the analysis involved factors with more than one degree of freedom in the numerator. Further, given a resolvable effect was significant both with and without a ROI interaction in a certain analysis, only the interaction involving ROI was resolved further.

## 6 Results

### 6.1 Behavioral data

The mean acceptability ratings for the stimuli, as well as the probe detection accuracy for the critical conditions, shown in Table 1, were calculated using the behavioral data collected during the experiment. Only those trials in which the acceptability judgement task following each trial was performed were included in the analysis. Further, the acceptability data presented here pertain only to the trials in which the participants performed the probe detection task correctly. Our behavioral results show that acceptability was very high for the acceptable conditions, HA and NA, and, albeit above chance, relatively low for the violation conditions, HV and NV. The difference in acceptability ratings between the acceptable and violation conditions was higher for sentences with human referents (92.18% vs. 59.76%) than for those with non-human referents (86.61% vs. 67.37). The overall accuracy was very high across all conditions. Owing to the fact that the reaction time data are not time-locked to the critical manipulation in the stimulus sentences, they are not reported here, but they are available from the corresponding author on request.

**Table 1:** Mean acceptability and accuracy ratings for the stimuli and probe detection for the critical conditions

| Condition | Acceptability % | SD | Accuracy % | SD |
| --- | --- | --- | --- | --- |
| Human-Acceptable (HA) | 92.18 | 12.27 | 95.18 | 6.37 |
| Human-Violation (HV) | 59.76 | 31.26 | 94.60 | 5.75 |
| Non-human-Acceptable (NA) | 86.61 | 15.67 | 95.62 | 6.61 |
| Non-human-Violation (NV) | 67.37 | 22.24 | 95.30 | 6.25 |

The statistical analysis of the behavioral data was performed by means of ANOVAs involving the within-subjects factors Humanness (HN) and Agreement type (AT), and the random factors participants (F1) and items (F2). Table 2 shows a summary of effects on the behavioral data collected during the experiment.

**Table 2:**
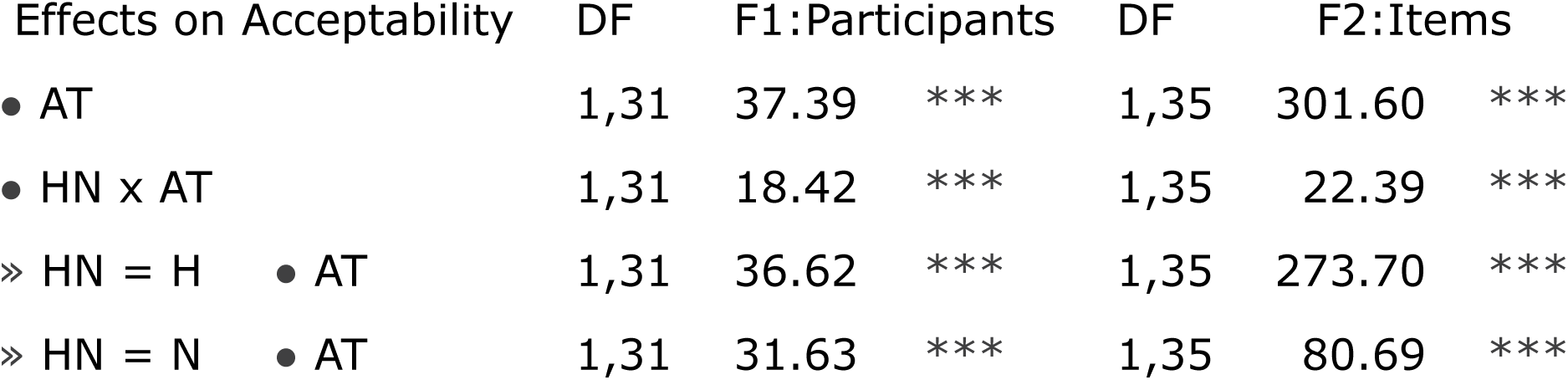
ANOVA of the behavioral data collected during the experiment. The following conventions are employed in the statistical tables: main effects and/or interactions that were at least marginally significant are reported; a factor or interaction following a bullet point means that factor or interaction was at least marginally significant; factorial resolutions are shown with an arrowhead followed by the factor and level; DF implies degrees of freedom; three stars beside F-values imply a significance of p ≤0.001; two stars imply p ≤0.01; a single start implies p ≤0.05; a single hollow circle implies a marginal (p ≤0.08) effect. Refer to Table 1 for the names of factors and levels used in this table.

| Effects on Acceptability | DF | F1:Participants | DF | F2:Items |
| --- | --- | --- | --- | --- |
| • AT | 1,31 | 37.39 *** | 1,35 | 301.60 *** |
| • HN x AT | 1,31 | 18.42 *** | 1,35 | 22.39 *** |
| » HN = H • AT | 1,31 | 36.62 *** | 1,35 | 273.70 *** |
| » HN = N • AT | 1,31 | 31.63 *** | 1,35 | 80.69 *** |

There was a main effect of Agreement on the acceptability in the analysis by participants, as well as in the analysis by items. The interaction Humanness x Agreement was significant in both analyses, which, when resolved for Humanness, showed an effect of Agreement for both human and non-human conditions. There were no effects on the probe detection accuracy.

### 6.2 ERP data

The ERPs at the adjective are shown in Figure 1 for the critical conditions. Three time-windows were chosen for analysis based on ERP components known to be relevant for agreement processing. Figure 2 shows the topographic map of the ERPs at the position of the adjective in the 300−500 ms, 500−700 ms and 900−1100 ms time-windows for the HV and NV conditions, after the effects for their corresponding acceptable condition has been subtracted. Table 3 shows a summary of all the effects that reached at least marginal significance at the position of the adjective in the selected time-windows.

**Figure 1:**
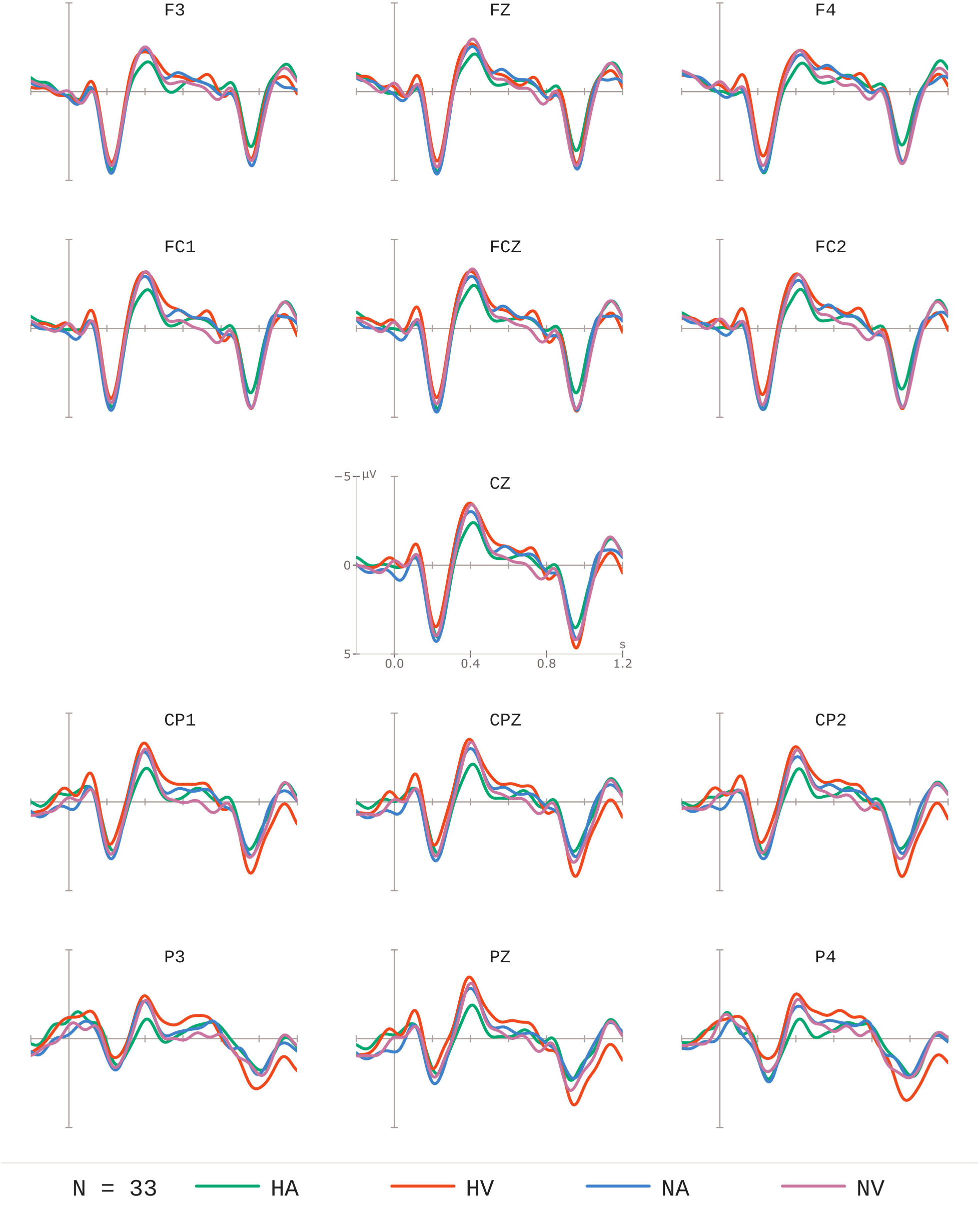
Grand-averaged ERPs time-locked to the position of the adjective for human and non-human violation conditions (red and purple lines) compared to their acceptable counterparts (green and blue lines).

**Figure 2:**
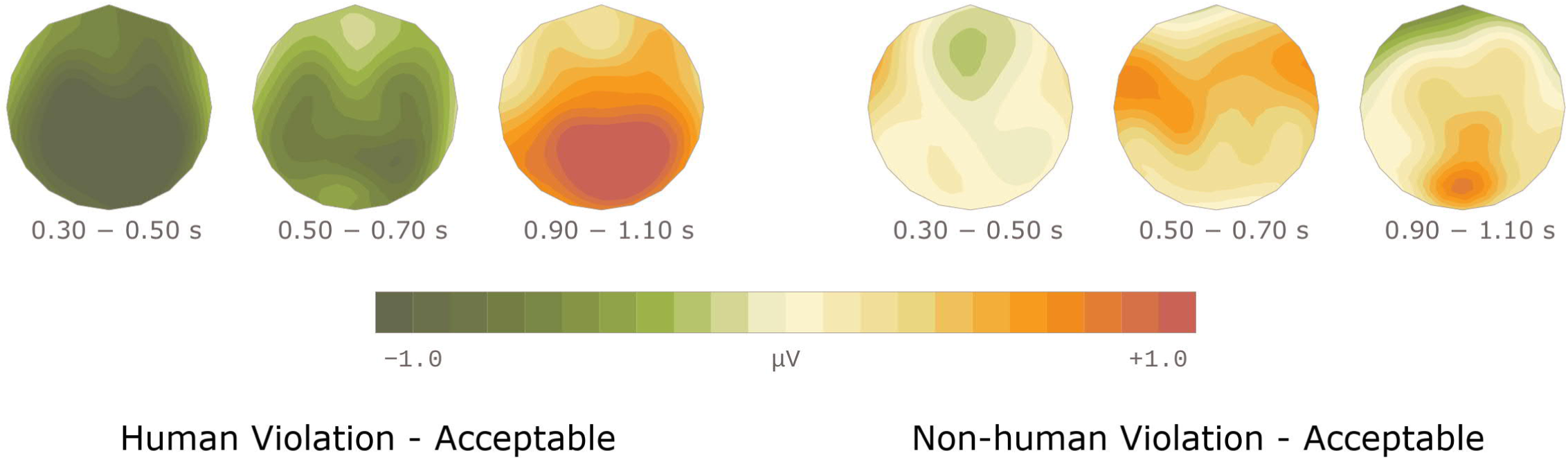
Topographic map of the ERPs at the position of the adjective in selected time-windows for human and non-human violation conditions, after the effects for the acceptable condition in each case has been subtracted.

**Table 3:** A summary of the ANOVA of ERP data at the Adjective position

| Effects in time-window 300–500 ms |  |  | DF | Lateral Regions |  | DF | Midline Regions |  |
| --- | --- | --- | --- | --- | --- | --- | --- | --- |
| • AT |  |  | 1,31 | 12.36 | *** | 1,31 | 15.07 | *** |
| • HN × AT |  |  | 1,31 | 6.77 | ** | 1,31 | 7.67 | ** |
| » HN = H | • AT |  | 1,31 | 19.03 | *** |  |  |  |
| • ROI × HN × AT |  |  |  |  |  | 5,155 | 3.52 | * |
| » ROI = CZ | • HN × AT |  |  |  |  | 1,31 | 5.52 | * |
|  | » HN = H | • AT |  |  |  | 1,31 | 18.86 | *** |
| » ROI = CPZ | • HN × AT |  |  |  |  | 1,31 | 12.81 | *** |
|  | » HN = H | • AT |  |  |  | 1,31 | 25.63 | *** |
| » ROI = PZ | • HN × AT |  |  |  |  | 1,31 | 14.22 | *** |
|  | » HN = H | • AT |  |  |  | 1,31 | 31.80 | *** |
| » ROI = POZ | • HN × AT |  |  |  |  | 1,31 | 11.08 | ** |
|  | » HN = H | • AT |  |  |  | 1,31 | 18.36 | *** |
| Effects in time-window 500–700 ms |  |  | DF | Lateral Regions |  | DF | Midline Regions |  |
| • ROI × HN |  |  | 3,93 | 3.85 | * |  |  |  |
| » ROI = LP | • HN |  | 1,31 | 7.78 | ** |  |  |  |
| • HN × AT |  |  | 1,31 | 8.30 | ** | 1,31 | 5.25 | * |
| » HN = H | • AT |  | 1,31 | 10.24 | ** | 1,31 | 7.09 | ** |
| » HN = N | • AT |  | 1,31 | 3.28 | . |  |  |  |
| Effects in time-window 900–1100 ms |  |  | DF | Lateral Regions |  | DF | Midline Regions |  |
| • ROI × HN |  |  | 3,93 | 14.41 | *** | 5,155 | 3.98 | * |
| » ROI = LA \|\| FZ | • HN |  | 1,31 | 6.50 | * | 1,31 | 7.17 | ** |
| » ROI = LP | • HN |  | 1,31 | 4.55 | * |  |  |  |
| » ROI = RP | • HN |  | 1,31 | 8.40 | ** |  |  |  |
| » ROI = CPZ | • HN |  |  |  |  | 1,31 | 3.46 | . |
| » ROI = PZ | • HN |  |  |  |  | 1,31 | 6.83 | * |
| » ROI = POZ | • HN |  |  |  |  | 1,31 | 3.18 | . |
| • AT |  |  | 1,31 | 7.48 | ** | 1,31 | 12.58 | *** |
| • ROI × AT |  |  | 3,93 | 5.81 | ** | 5,155 | 6.76 | ** |
| » ROI = LP \|\| FCZ | • AT |  | 1,31 | 7.82 | ** | 1,31 | 4.60 | * |
| » ROI = CZ | • AT |  |  |  |  | 1,31 | 5.58 | * |
| » ROI = RP \|\| CPZ | • AT |  | 1,31 | 13.71 | *** | 1,31 | 13.04 | *** |
| » ROI = PZ | • AT |  |  |  |  | 1,31 | 14.44 | *** |
| » ROI = POZ | • AT |  |  |  |  | 1,31 | 14.62 | *** |
| • HN × AT |  |  | 1,31 | 4.93 | * |  |  |  |
| » HN = H | • AT |  | 1,31 | 17.50 | *** |  |  |  |

| Effects in time-window 300–500 ms |  |  |  | DF | Lateral Regions |  | DF | Midline Regions |  |
| --- | --- | --- | --- | --- | --- | --- | --- | --- | --- |
| ● AT |  |  |  | 1,31 | 12.36 | *** | 1,31 | 15.07 | *** |
| ● HN x AT |  |  |  | 1,31 | 6.77 | ** | 1,31 | 7.67 | ** |
| » HN = H | ● AT |  |  | 1,31 | 19.03 | *** |  |  |  |
| ● ROI x HN x AT |  |  |  |  |  |  | 5,155 | 3.52 | * |
| » ROI = CZ | ● HN x AT |  |  |  |  |  | 1,31 | 5.52 | * |
|  |  | » HN = H | ● AT |  |  |  | 1,31 | 18.86 | *** |
| » ROI = CPZ | ● HN x AT |  |  |  |  |  | 1,31 | 12.81 | *** |
|  |  | » HN = H | ● AT |  |  |  | 1,31 | 25.63 | *** |
| » ROI = PZ | ● HN x AT |  |  |  |  |  | 1,31 | 14.22 | *** |
|  |  | » HN = H | ● AT |  |  |  | 1,31 | 31.80 | *** |
| » ROI = POZ | ● HN x AT |  |  |  |  |  | 1,31 | 11.08 | ** |
|  |  | » HN = H | ● AT |  |  |  | 1,31 | 18.36 | *** |
| Effects in time-window 500–700 ms |  |  |  | DF | Lateral Regions |  | DF | Midline Regions |  |
| ● ROI x HN |  |  |  | 3,93 | 3.85 | * |  |  |  |
| » ROI = LP | ● HN |  |  | 1,31 | 7.78 | ** |  |  |  |
| ● HN x AT |  |  |  | 1,31 | 8.30 | ** | 1,31 | 5.25 | * |
| » HN = H | ● AT |  |  | 1,31 | 10.24 | ** | 1,31 | 7.09 | ** |
| » HN = N | ● AT |  |  | 1,31 | 3.28 | ◦ |  |  |  |
| Effects in time-window 900–1100 ms |  |  |  | DF | Lateral Regions |  | DF | Midline Regions |  |
| ● ROI x HN |  |  |  | 3,93 | 14.41 | *** | 5,155 | 3.98 | * |
| » ROI = LA \|\| FZ | ● HN |  |  | 1,31 | 6.50 | * | 1,31 | 7.17 | ** |
| » ROI = LP | ● HN |  |  | 1,31 | 4.55 | * |  |  |  |
| » ROI = RP | ● HN |  |  | 1,31 | 8.40 | ** |  |  |  |
| » ROI = CPZ | ● HN |  |  |  |  |  | 1,31 | 3.46 | ◦ |
| » ROI = PZ | ● HN |  |  |  |  |  | 1,31 | 6.83 | * |
| » ROI = POZ | ● HN |  |  |  |  |  | 1,31 | 3.18 | ◦ |
| ● AT |  |  |  | 1,31 | 7.48 | ** | 1,31 | 12.58 | *** |
| ● ROI x AT |  |  |  | 3,93 | 5.81 | ** | 5,155 | 6.76 | ** |
| » ROI = LP \|\| FCZ | ● AT |  |  | 1,31 | 7.82 | ** | 1,31 | 4.60 | * |
| » ROI = CZ | ● AT |  |  |  |  |  | 1,31 | 5.58 | * |
| » ROI = RP \|\| CPZ | ● AT |  |  | 1,31 | 13.71 | *** | 1,31 | 13.04 | *** |
| » ROI = PZ | ● AT |  |  |  |  |  | 1,31 | 14.44 | *** |
| » ROI = POZ | ● AT |  |  |  |  |  | 1,31 | 14.62 | *** |
| ● HN x AT |  |  |  | 1,31 | 4.93 | * |  |  |  |
| » HN = H | ● AT |  |  | 1,31 | 17.50 | *** |  |  |  |

#### 6.2.1 Time-window 300−500 ms

The predominant effect in the 300 – 500 ms time-window is the negativity for the violation condition as opposed to the acceptable condition, but only when the subject noun was human. There is virtually no difference between the acceptable and violation conditions when the subject was non-human. There was a main effect of Agreement in both the lateral and the midline regions. The interaction Humanness x Agreement was likewise significant in all the regions. Resolving this for the levels of Humanness showed an effect of Agreement in the lateral regions. In the midline regions, the three-way interaction ROI x Humanness x Agreement was significant, which when resolved for the levels of midline ROIs showed that the interaction Humanness x Agreement was significant in the central, centroparietal, parietal, and parietooccipital midline regions. Resolving this further for the levels of Humanness revealed a simple effect of Agreement in all the regions mentioned.

#### 6.2.2 Time-window 500−700 ms

The negativity in the previous time-window for the violations as opposed to acceptable sentences involving human subject nouns continues to remain significant in the 500 – 700 ms time-window. There were no main effects. The interaction ROI x Humanness was significant in the lateral regions. Resolving this for the individual ROIs showed a significant effect of Humanness in the left-posterior region. The interaction Humanness x Agreement was significant in the lateral as well as midline regions, which when resolved showed a significant effect of Agreement in all regions when the subject noun was human. There was a marginal effect of Agreement in the lateral electrode regions when the subject was non-human.

#### 6.2.3 Time-window 900-1100 ms

A general effect of Agreement begins to emerge in this time-window, which is nevertheless qualified by a Humanness interaction, whereby the violation conditions elicited more positive-going ERPs as opposed to the acceptable conditions for human noun conditions. There is a main effect of Agreement in all electrode regions. The interaction ROI x Humanness reached significance in all the regions, which when resolved for the individual ROIs revealed an effect of Humanness in all left-anterior, left-posterior and right-posterior lateral electrode regions and the frontal and parietal midline regions. The effect of Humanness was marginal in the centroparietal and parietooccipital midline regions. The interaction ROI x Agreement was significant in all the regions, which was resolved for the individual ROIs. This showed an effect of Agreement in the left-posterior and right-posterior lateral regions, as well as in all the midline regions with the exception of the frontal midline region. The interaction Humanness x Agreement was significant in the lateral regions alone, which when resolved showed a simple effect of Agreement for human noun conditions.

## 7 General discussion

ERPs at the position of the adjective showed that violations of agreement in the context of a human referent (HV) evoked a negativity effect followed by a late-positivity effect, as opposed to their acceptable counterparts (HA). The negativity effect was significant in a broad time-window spanning from 300 ms until about 700 ms after the onset of the adjective. No such effect ensued in the non-human violation (NV) conditions. Based on the onset latency and topographic distribution of these effects, we interpret the negativity as an instance of an N400 effect, and the late-positivity as an instance of a P600 effect.

### 7.1 The biphasic response

Our finding of a biphasic N400-P600 effect is in line with a number of studies in other languages involving agreement violations of various types of dependencies (i.e., subject-verb, adjective-noun, article-noun etc.). These include: number violations in Basque (Zawiszewski et al., 2016); gender violations in Dutch (Hagoort et al., 1993), Hebrew (Deutsch & Bentin, 2001; with an animacy interaction), Spanish (Guajardo & Wicha, 2014); and person violations in Basque (Mancini et al., 2019; Zawiszewski & Friederici, 2009; Zawiszewski et al., 2016) and Spanish (Mancini et al., 2011). Agreement violations are traditionally viewed as formal morphosyntactic rule violations. These violations have been commonly associated with concomitant LAN effects interpreted as indicative of the detection of the morphosyntactic violation (Bornkessel & Schlesewsky, 2006; Friederici, 2002; Münte, Matzke, & Johannes, 1997; Münte, Szentkuti, et al., 1997). However, Molinaro et al. (2011) note that the nature of the complexities involved in morphological decomposition for feature identification may determine whether a LAN or an N400 ensues. Indeed, Choudhary, Schlesewsky, Roehm, & Bornkessel-Schlesewsky (2009) argue that the dissociation between a LAN and an N400 results from whether an interpretively relevant cue is violated, in which case an N400 arises; or alternatively whether the violation involves a cue that is irrelevant for interpretation, in which case a LAN ensues. In view of this, the absence of a LAN effect in our study is not surprising, given that agreement computation in Arabic is not simply formal but depends on specific syntactic properties of the construction involved, such as word-order and whether or not the subject is overt, as well as on properties at the syntax-semantic interface such as humanness/animacy (for a detailed discussion, see pre-print of Muralikrishnan & Idrissi (2019), article in review). Agreement in Arabic crucially depends on number and gender features, both of which are highly relevant cues for interpretation. Our finding of a biphasic N400 (rather than LAN) -P600 effect for human violations is therefore not surprising.

The N400 effect we observed may also be taken to reflect the cognitive routines involved in the process of verifying, at the lexical-semantic level, that the morphological properties of the adjective match the semantic and/or syntactic properties of the preceding plural noun. The mismatch in the case of human violations leads to a larger N400 (Friederici, 2011; Molinaro et al., 2011), because the morphosyntactic information encoded on the adjective (i.e., feminine singular morphology) fails to map onto the gender and number specifications of the noun (i.e., masculine plural semantics and/or morphology) during the construction of sentence meaning.

As in several studies cited above involving agreement violations, the P600 effect in our study can be plausibly interpreted as reflecting a well-formedness check (Bornkessel & Schlesewsky, 2006; Friederici, 2011; Mancini et al., 2019) or syntactic integration costs (Barber et al., 2004; Kaan et al., 2000). Late positivities are said to arise at the integration/interpretation stage and are generally associated with structural processing, and specifically, with reanalysis routines activated in response to ungrammatical, ambiguous or complex information. P600-type late positive effects can also be interpreted as signaling domain-general conflict-monitoring processes which are not limited to language (and syntactic analysis) per se (Mancini et al., 2019; van de Meerendonk et al., 2009). In terms of language processing, the P600 effect, under this view, arises when a strong expectation conflicts with the actual input. Such a conflict initiates procedures of reanalysis/repair upon detection of the errors in the input. This may indeed be what happens during the processing of the human violations in the current study. Specifically, an adjective is expected to fully agree in number and gender with a preceding masculine plural human noun. But, when it is marked as feminine singular, instead, a conflict arises between the expectation and the actual input, and procedures of reanalysis/repair are then initiated. By contrast, a similar response does not ensue when DA is violated, since DA is less strict than FA, and the expectation that it obtains is therefore less strong. The resulting conflict is therefore never serious enough to trigger noticeable repair processes. The fact that the repair process takes place in the later (800 – 1100 ms) time window may be said to further support this view, and may additionally be indicative of suppression by the parser of the repair routines due to the prevalence of DA in the language (there are far fewer human nouns than animal and inanimate nouns) and/or the fact that DA and FA are not completely mutually exclusive in Spoken Arabic.

The relatively long latency of the N400 reported in our study may be due to the increased processing (and memory) cost at the syntax-semantics interface (Fiebach, Schlesewsky, & Friederici, 2002) resulting from the complex interaction of morphology and humanness (animacy), in Arabic, as well as from the possibility which speakers have to assign the majority of plural nouns in their spoken varieties both a plural and collective/singulative semantics (see Dali & Mathieu (2016)).

Clearly, our results illustrate the central role animacy/humanness plays in grammatical agreement in Arabic. Animacy has been shown to be a highly salient feature in typologically diverse languages such as Mandarin Chinese (Philipp, Bornkessel-Schlesewsky, Bisang, & Schlesewsky, 2008), Polish (Szewczyk & Schriefers, 2011, 2013), and Tamil (Muralikrishnan, Schlesewsky, & Bornkessel-Schlesewsky, 2015). Therefore, it is no surprise that humanness, which tops the animacy hierarchy (Comrie, 1989; Silverstein, 1976) is a highly salient feature in Arabic that interacts with plurality and gender. Recall that Standard Arabic requires that non-human (animate and inanimate) masculine plural nouns be treated as 3rd singular feminine nouns as far as agreement is concerned, such that the canonical rule (i.e., matching number and gender feature agreement between the adjective and the noun) would constitute a violation. By contrast, human masculine plural nouns do require the canonical agreement pattern (i.e., full feature matching). This results in an overall distribution in Standard Arabic where DA (the exception) is as good and as frequent as (if not more than) canonical agreement (i.e., FA), because non-human animate and inanimate referents outnumber human referents in the language. This in turn leads to a situation in which a violation at the position of the adjective following a human noun (i.e., DA instead of the required FA) would not constitute an outright violation, because DA is as frequent in the language as FA for this type of noun. Nevertheless, given full lexical access of the preceding noun would have indicated it to be human, this property renders DA a violation. In other words, the animacy cue is more salient and crucial than the agreement feature cues in this regard, and in fact the two types of cues contradict each other as far as the correct agreement paradigm is concerned. Therefore, the processing system has to resolve this conflict in order to come up with an evaluation about whether a violation is involved. The complexity involved in this process contributes to the latency of the negativity effect. This complexity is further enhanced by the agreement conventions of the spoken variety, which for human nouns coincide with the agreement required by the animacy cue but are significantly variable in the case of nonhuman nouns. The variation in the behavioral acceptability judgements for violations involving human referents attests to this fact.

Converging evidence for such a processing complexity-based interpretation comes from a study on Hindi agreement involving animacy by Bhattamishra, Muralikrishnan and Choudhary (submitted), who also report a late-latency negativity similar to the one we found in our study. Further, in a recent article, Bornkessel-Schlesewsky & Schlesewsky (2019) argue for a neurobiologically plausible model, where they posit that all the language-related negativities form a family of functionally related rather than distinct negativities, and that their different latencies reflect the level of complexity involved in processing the stimulus.

A non-trivial aspect to consider in explaining our results is also the availability of alternative agreement patterns in the grammar(s) of Arabic diglossic speakers. DA with human nouns should show a greater mismatch since human nouns show more stable gender and number specifications than non-human nouns. Violations of FA with human nouns thus leads to a neurophysiological response that is typical for morphosyntactic violations. By contrast, FA with non-human nouns (which is supposed to be a violation) should not matter as much since non-human plurals show underlying representations of so-called ‘hybrid’ nouns (see Landau, 2016), and as such may trigger either semantic agreement (which coincides with DA) or morphological/syntactic agreement (which coincides with FA). The tolerance of the mismatch (between semantic and morphological features) which we observed in favor of the match (between syntactic and morphological features) (i.e., of DA) must thus arise as a result of the architecture of the Arabic speakers’ diglossic grammar. As mentioned earlier, non-human masculine plural nouns require DA in Standard Arabic, while DA is not mandatory in the spoken varieties, in which FA and DA tend to be both acceptable.

Given that our results show that human plural nouns trigger FA, and any human plural noun-adjective construction that does not adhere to FA is ungrammatical, we argue that our first hypothesis is confirmed, with the critical role of humanness in the Arabic agreement patterns being confirmed. Obviously, adherence to DA is not as compulsory for non-human plural nouns, a state of affairs most likely caused by interference from Spoken Arabic where nonhuman plural nouns allow FA in addition to DA. This supports our second hypothesis regarding the effect of diglossia, a point we further discuss next.

### 7.2 Diglossic grammar

As mentioned above, the behavioral and ERP results reported in this study show that the processing of Standard Arabic structures is influenced by the representations and rules of Spoken Arabic. This should be expected if we assume that in terms of underlying lexical representations, Standard Arabic and Spoken Arabic share the same lexical repertoire (see the results of a recent fMRI study by Abou-Ghazaleh, Khateb, & Nevat, 2018). This entails that the lexical representations of Standard and Spoken Arabic (plural) nouns could overlap or even be identical. Additionally, if we assume that the representations of Spoken Arabic dominate (probably due to the level of competence, extent of usage, and the mother tongue vs. second language status), we could explain why the highly hybrid nature of non-human plural nouns in Spoken Arabic modulates how they are processed in Standard Arabic.

That HV elicited the biphasic negativity-late positivity effect, while NV did not, indicates that violations of DA, unlike violations of FA, are not processed as outright violations. We ascribe this to the status of the DA pattern in Spoken Arabic. As mentioned earlier, non-human masculine plural nouns require DA in Standard Arabic; whereas DA is not mandatory in the spoken varieties; rather, both FA and DA are allowed. Violations of DA are thus not processed as anomalies, and indeed they pattern with their acceptable counterpart sentences. This makes sense under a convergence theory of the lexical/semantic representations and syntactic competence of our participants, whereby words share similar or the same lexical features, and the syntactic rules of Standard Arabic and Spoken Arabic coexist within one larger system (Ameel, Malt, Storms, & Van Assche, 2009; Green, 2003; Green, Crinion, & Price, 2006; White, Malt, & Storms, 2017).

Within a straightforward syntactic analysis of FA and DA, one can posit that human plurals in Standard Arabic bear the right features for gender and number at an underlying level, and these features are checked as expected during the interpretation of the sentence. As for non-human nouns, they are underlyingly specified as feminine and singular, which explain the surface DA pattern (Dali & Mathieu, 2016). Like the masculine and plural in FA, the features feminine and singular in DA are checked during the computation of sentence structure and meaning. Thus, disagreement triggered by non-human referents does not actually constitute failure of agreement or disagreement. When human, the trigger nouns are most likely to bear the expected natural gender (i.e., [+masculine]), and, by virtue of their position at the top of Individuation Hierarchy (Audring, 2008; Brustad, 2000; Sasse, 1993) (a version of the Animacy hierarchy), the expected number feature [+plural]. Barlow (1988) argues that high agency, animacy, familiarity, and clear definition of individual entities (specificity and clear boundaries between them) define individuality and how a group may be treated as a set of individuals (and therefore, as a plural) or as a unit/group (and therefore, as a singular). Human nouns in Standard Arabic are then systematically assigned a distributive interpretation and are treated as plurals. By contrast, nonhuman animal and inanimate referents, being respectively lower and lowest on the Individuation Hierarchy, are systematically assigned a collective or singulative interpretation, with which the feminine happens to be associated in the language (Dali & Mathieu, 2016; Fassi Fehri, 2016).

In Spoken Arabic, broken plurals are *highly* ‘hybrid’ nouns in that agreement can target either their syntactic (i.e., [+masculine] and [+plural]) or (lexically specified) semantic features (i.e., [-plural, +group]) (Landau, 2016). Feminine agreement on broken plurals is the realization of the number feature [+group] (Dali & Mathieu, 2016). Variation is thus expected in Spoken Arabic. Cowell (1964:423) observes that in Damascus Arabic, DA is assigned to inanimate plurals and some animate plurals and collectives, when the meaning of “collectivity or generality is emphasized rather than heterogeneity or particularity.”

Compared to their counterparts in Standard Arabic, the underlying representations of plural nouns are slightly different in Spoken Arabic allowing for alternative gender and number feature specifications. Human and inanimate referents are mostly like in Standard Arabic, whereas animal referents allow both distributive and singulative interpretations, the selection of which is determined by various semantic, pragmatic and discourse factors. This explains both the patterns reported by Belnap (1993) as well as our behavioral and ERP data.

Of course, if only the Standard Arabic system were consulted during the processing of the critical sentences in the present study, one would expect nonhuman violations to elicit the same behavioral and ERP patterns as human violations when DA is violated. Yet, this does not happen because the featural conflict (in the sense of not abiding by the expected pattern) is resolved under pressure from the Spoken Arabic grammar, in which masculine animal plurals can also be interpreted, and therefore represented, as [+masculine, +plural]. This is reflected in the absence of any significant difference between nonhuman violations and nonhuman acceptable structures in terms of the ERPs they trigger throughout all the critical time windows.

Clearly then, the processing system of our participants must consult a linguistic system in which two sub-systems, Standard Arabic and Spoken Arabic, coexist and can be in competition when their representations and rules diverge. In this diglossic system, while the diverging patterns compete, this competition is resolved as each subsystem yields in to the other. Since neurophysiological research on Arabic is still in its infancy, however, future research is needed to explore various aspects of grammatical agreement and the impact of diglossia on language processing. For example, one should explore whether a design including inanimate plurals would elicit a parametric difference between human violations, nonhuman animate violations, and nonhuman inanimate violations. Note that inanimate plural nouns fall on the other end of the continuum as they show more stable lexical representations than nonhuman animate ones. The impact of diglossia on other aspects of language processing in the language also needs to be examined further and more systematically. Finally, it would be interesting to explore whether the same patterns observed in written sentence comprehension would be observed in auditory sentence comprehension, and whether the variety in which the auditory sentences appear, Standard or Spoken Arabic, would make a difference.

### 8 Conclusions and future work

This paper set out to investigate the processing of full and deflected agreement in Standard Arabic. Given that Arabic is a widely spoken but highly understudied language, and in view of the diglossic language situation of Arabic speakers, our study constitutes an important contribution to the literature on online sentence processing. The results reported here shed more light on the neurobiological foundations of language and provide crucial insights into the processing of a non-canonical grammatical agreement pattern. Our results add further support for the relationship between processing complexity and the latency of negativity effects, and for the importance of features at the syntax-semantics interface such as animacy in language comprehension. We show that the presence of optional grammars shapes the dynamics of language processing, as illustrated by the differential treatment of human violations and nonhuman violations of agreement, and the prolonged negativity and the latency of P600. Our findings suggest that the diglossic brain has at its disposal a ‘converging’ grammar where conflicting patterns are ‘attenuated’ into a compromise system. The manifestations of this compromise system/grammar can been seen throughout early and later stages of written sentence comprehension.

## 9 Acknowledgments

This work was supported by the Qatar National Research Fund; the National Priorities Research Program grant number [NPRP 7-427-6-011].

## Footnotes

1 The following abbreviations are used: m. = masculine; sg. = singular; pl. = plural; f. = feminine; and 1, 2 and 3 = first, second and third person, respectively. Case endings are not indicated, and the feminine suffix is transcribed as /-at/ although the consonant is not always pronounced.

2 We use Corbett’s (2006) distinction between ‘syntactic’ and ‘semantic’ agreement. Syntactic agreement refers to cases where the morphology of the target is consistent with the morphosyntactic properties of the controller, while semantic agreement refers to cases where the target echoes the semantics properties of the controller.

## References

Abou-Ghazaleh, A., Khateb, A., & Nevat, M. (2018). Lexical Competition between Spoken and Literary Arabic: A New Look into the Neural Basis of Diglossia Using fMRI. Neuroscience, 393, 83–96. https://doi.org/10.1016/j.neuroscience.2018.09.045

Alday, P. M., Schlesewsky, M., & Bornkessel-Schlesewsky, I. (2017). Electrophysiology Reveals the Neural Dynamics of Naturalistic Auditory Language Processing: Event-Related Potentials Reflect Continuous Model Updates. Eneuro, 4(6), ENEURO.0311-16.2017. https://doi.org/10.1523/ENEURO.0311-16.2017

Ambros, A. A. (1977). Damascus Arabic. Afroasiatic Dialects: Semitic, 3, Malibu, California: Undena Publications.

Ameel, E., Malt, B. C., Storms, G., & Van Assche, F. (2009). Semantic convergence in the bilingual lexicon. Journal of Memory and Language, 60(2), 270–290. https://doi.org/10.1016/j.jml.2008.10.001

Angrilli, A., Penolazzi, B., Vespignani, F., De Vincenzi, M., Job, R., Ciccarelli, L., … Stegagno, L. (2002). Cortical brain responses to semantic incongruity and syntactic violation in Italian language: An event-related potential study. Neuroscience Letters, 322(1), 5–8. https://doi.org/10.1016/S0304-3940(01)02528-9

Aoun, E. J., Benmamoun, E., & Choueiri, L. (2010). The Syntax of Arabic. Cambridge, UK: Cambridge University Press.

Audring, J. (2008). Gender assignment and gender agreement: Evidence from pronominal gender languages. Morphology, 18(2), 93–116. https://doi.org/10.1007/s11525-009-9124-y

Barber, H., & Carreiras, M. (2003). Integrating Gender and Number Information in Spanish Word Pairs: An Erp Study. Cortex, 39(3), 465–482. https://doi.org/10.1016/S0010-9452(08)70259-4

Barber, H., & Carreiras, M. (2005). Grammatical gender and number agreement in Spanish: An ERP comparison. Journal of Cognitive Neuroscience, 17(1), 137–153.

Barber, H., Salillas, E., & Carreiras, M. (2004). Gender or genders agreement. On-Line Study of Sentence Comprehension, 309–328.

Barlow, M. (1988). A Situated Theory of Agreement (Ph.D. Dissertation). Stanford University.

Belnap, R. K. (1993). The meaning of deflected/strict agreement variation in Cairene Arabic. In Perspectives on Arabic Linguistics 5 (pp. 97–118). Amsterdam/Philadelphia: John Benjamins Publishing Company.

Belnap, R. K., & Haeri, N. (1997). Structuralist Studies in Arabic Linguistics: Charles A. Ferguson’s Papers, 1854–1994. Leiden: Brill.

Bettega, S. (2017). Agreement with Plural Controllers in Omani Arabic: Preliminary remarks. In S. Bettega & F. Gasparini (Eds.), Linguistic Studies in the Arabian Gulf (pp. 153–174). Torino: Dipartimento di Lingue e Letterature straniere e Culture moderne – Università di Torino.

Bornkessel, I., & Schlesewsky, M. (2006). The extended argument dependency model: A neurocognitive approach to sentence comprehension across languages. Psychological Review, 113(4), 787–821. https://doi.org/10.1037/0033-295X.113.4.787

Bornkessel-Schlesewsky, I., & Schlesewsky, M. (2008). An alternative perspective on “semantic P600” effects in language comprehension. Brain Research Reviews, 59(1), 55–73. https://doi.org/10.1016/j.brainresrev.2008.05.003

Bornkessel-Schlesewsky, I., & Schlesewsky, M. (2019). Toward a Neurobiologically Plausible Model of Language-Related, Negative Event-Related Potentials. Frontiers in Psychology, 10. https://doi.org/10.3389/fpsyg.2019.00298

Brustad, K. (2000). The syntax of spoken Arabic. Georgetown University Press.

Choudhary, K. K., Schlesewsky, M., Roehm, D., & Bornkessel-Schlesewsky, I. (2009). The N400 as a correlate of interpretively relevant linguistic rules: Evidence from Hindi. Neuropsychologia, 47(13), 3012–3022. https://doi.org/10.1016/j.neuropsychologia.2009.05.009

Comrie, B. (1989). Language Universals and Linguistic Typology: Syntax and Morphology (2nd Edition). Berlin, Boston: De Gruyter Mouton.

Corbett, G. G. (2006). Agreement. Cambridge: Cambridge University Press.

Cowell, M. W. (1964). A Reference Grammar of Syrian Arabic, based on the dialect of Damascus. (=Arabic series, 4.). Washington, D.C.: Georgetown University, Institute of Languages and Linguistics.

Dali, M., & Mathieu, E. (2016). Les pluriels internes féminins de l’arabe tunisien. Lingvisticæ Investigationes, 39(2), 253–271.

D’Anna, L. (2017). Agreement with plural controllers in Fezzānī Arabic. *Folia Orientalia*, LIV, 23.

Deutsch, A., & Bentin, S. (2001). Syntactic and Semantic Factors in Processing Gender Agreement in Hebrew: Evidence from ERPs and Eye Movements. Journal of Memory and Language, 45(2), 200–224. https://doi.org/10.1006/jmla.2000.2768

Díaz, B., Sebastián-Gallés, N., Erdocia, K., Mueller, J. L., & Laka, I. (2011). On the cross-linguistic validity of electrophysiological correlates of morphosyntactic processing: A study of case and agreement violations in Basque. Journal of Neurolinguistics, 24(3), 357–373. https://doi.org/10.1016/j.jneuroling.2010.12.003

Fassi Fehri, A. (2016). Semantic Gender Diversity and Its Architecture in the Grammar of Arabic. Brill’s Journal of Afroasiatic Languages and Linguistics, 8(1), 154–199. https://doi.org/10.1163/18776930-00801007

Fassi-Fehri, A. (1981). Théorie lexicale-fonctionelle, controle et accord en arabe moderne. Arabica, 28, 299–332.

Ferguson, C. A. (1989). Grammatical Agreement in Classical Arabic and the Modern Dialects: A Response to Versteegh’s Pidginization Hypothesis. Al-’Arabiyya, 22(1/2), 5–17.

Fiebach, C. J., Schlesewsky, M., & Friederici, A. D. (2002). Separating syntactic memory costs and syntactic integration costs during parsing: The processing of German WH-questions. Journal of Memory and Language, 47(2), 250–272. https://doi.org/10.1016/S0749-596X(02)00004-9

Friederici, A. D. (1995). The Time Course of Syntactic Activation During Language Processing: A Model Based on Neuropsychological and Neurophysiological Data. Brain and Language, 50(3), 259–281. https://doi.org/10.1006/brln.1995.1048

Friederici, A. D. (2002). Towards a neural basis of auditory sentence processing. Trends in Cognitive Sciences, 6(2), 78–84. https://doi.org/10.1016/S1364-6613(00)01839-8

Friederici, A. D. (2004). Event-related brain potential studies in language. Current Neurology and Neuroscience Reports, 4(6), 466–470. https://doi.org/10.1007/s11910-004-0070-0

Friederici, A. D. (2011). The Brain Basis of Language Processing: From Structure to Function. Physiological Reviews, 91(4), 1357–1392. https://doi.org/10.1152/physrev.00006.2011

Green, D. W. (2003). Neural basis of lexicon and grammar in L2 acquisition: The convergence hypothesis. In R. van Hout, A. Hulk, F. Kuiken, & R. Towell (Eds.), The lexicon-syntax interface in second language acquisition. (pp. 197–218). Retrieved from http://discovery.ucl.ac.uk/164901/

Green, D. W., Crinion, J., & Price, C. J. (2006). Convergence, degeneracy and control. Language Learning, 56(S1), 99–125. Retrieved from https://www.ncbi.nlm.nih.gov/pmc/articles/PMC2241761/

Guajardo, L. F., & Wicha, N. Y. Y. (2014). Morphosyntax can modulate the N400 component: Event related potentials to gender-marked post-nominal adjectives. NeuroImage, 91, 262–272. https://doi.org/10.1016/j.neuroimage.2013.09.077

Hagoort, P. (2003). Interplay between syntax and semantics during sentence comprehension: ERP effects of combining syntactic and semantic violations. Journal of Cognitive Neuroscience, 15(6), 883–899.

Hagoort, P., Brown, C., & Groothusen, J. (1993). The syntactic positive shift (SPS) as an ERP measure of syntactic processing. Language and Cognitive Processes, 8(4), 439–483.

Hinojosa, J., Martín-Loeches, M., Casado, P., Muñoz, F., & Rubia, F. (2003). Similarities and differences between phrase structure and morphosyntactic violations in Spanish: An event-related potentials study. Language and Cognitive Processes, 18(2), 113–142. https://doi.org/10.1080/01690960143000489

Huynh, H., & Feldt, L. S. (1970). Conditions Under Which Mean Square Ratios in Repeated Measurements Designs Have Exact F-Distributions. Journal of the American Statistical Association, 65(332), 1582. https://doi.org/10.2307/2284340

Kaan, E., Harris, A., Gibson, E., & Holcomb, P. (2000). The P600 as an index of syntactic integration difficulty. Language and Cognitive Processes, 15(2), 159–201. https://doi.org/10.1080/016909600386084

Kutas, M., & Hillyard, S. A. (1983). Event-related brain potentials to grammatical errors and semantic anomalies. Memory & Cognition, 11(5), 539–550.

Landau, I. (2016). DP-internal semantic agreement: A configurational analysis. Natural Language & Linguistic Theory, 34(3), 975–1020.

Lau, E. F., Phillips, C., & Poeppel, D. (2008). A cortical network for semantics: (De)constructing the N400. Nature Reviews Neuroscience, 9(12), 920–933. https://doi.org/10.1038/nrn2532

Leinonen, A., Brattico, P., Järvenpää, M., & Krause, C. M. (2008). Event-related potential (ERP) responses to violations of inflectional and derivational rules of Finnish. Brain Research, 1218, 181–193. https://doi.org/10.1016/j.brainres.2008.04.049

Mancini, S., Massol, S., Duñabeitia, J. A., Carreiras, M., & Molinaro, N. (2018). Agreement and illusion of disagreement: An ERP study on Basque. Cortex, S001094521830296X. https://doi.org/10.1016/j.cortex.2018.08.036

Mancini, S., Massol, S., Duñabeitia, J. A., Carreiras, M., & Molinaro, N. (2019). Agreement and illusion of disagreement: An ERP study on Basque. Cortex, 116, 154–167. https://doi.org/10.1016/j.cortex.2018.08.036

Mancini, S., Molinaro, N., Rizzi, L., & Carreiras, M. (2011). When persons disagree: An ERP study of Unagreement in Spanish: Unagreement in Spanish. Psychophysiology, 48(10), 1361–1371. https://doi.org/10.1111/j.1469-8986.2011.01212.x

Martin-Loeches, M., Nigbur, R., Casado, P., Hohlfeld, A., & Sommer, W. (2006). Semantics prevalence over syntax during sentence processing: A brain potential study of nouneadjective agreement in Spanish. Brain Research, 1093(1), 178–189.

Melebari, A. (2017). The interaction of Animacy and Morphosyntax in Arabic. Stony Brook University.

Molinaro, N., Barber, H. A., & Carreiras, M. (2011). Grammatical agreement processing in reading: ERP findings and future directions. Cortex, 47(8), 908–930. https://doi.org/10.1016/j.cortex.2011.02.019

Molinaro, N., Vespignani, F., & Job, R. (2008). A deeper reanalysis of a superficial feature: An ERP study on agreement violations. Brain Research, 1228, 161–176. https://doi.org/10.1016/j.brainres.2008.06.064

Münte, T. F., Matzke, M., & Johannes, S. (1997). Brain activity associated with syntactic incongruencies in words and pseudo-words. Journal of Cognitive Neuroscience, 9(3), 318–329. https://doi.org/10.1162/jocn.1997.9.3.318

Münte, T. F., Szentkuti, A., Wieringa, B. M., Matzke, M., & Johannes, S. (1997). Human brain potentials to reading syntactic errors in sentences of different complexity. Neuroscience Letters, 235(3), 105–108. https://doi.org/10.1016/s0304-3940(97)00719-2

Muralikrishnan, R., & Idrissi, A. (2019). Cognitive salience of agreement features modulates language comprehension. BioRxiv, 671834. https://doi.org/10.1101/671834

Muralikrishnan, R., Schlesewsky, M., & Bornkessel-Schlesewsky, I. (2015). Animacy-based predictions in language comprehension are robust: Contextual cues modulate but do not nullify them. Brain Research, 1608, 108–137. https://doi.org/10.1016/j.brainres.2014.11.046

Nevins, A., Dillon, B., Malhotra, S., & Phillips, C. (2007). The role of feature-number and feature-type in processing Hindi verb agreement violations. Brain Research, 1164, 81–94. https://doi.org/10.1016/j.brainres.2007.05.058

Nichols, J., & Bickel, B. (2013). Locus of Marking: Whole-language Typology. In M. S. Dryer & M. Haspelmath (Eds.), The World Atlas of Language Structures Online. Retrieved from https://wals.info/chapter/25

O’Rourke, P. L., & Van Petten, C. (2011). Morphological agreement at a distance: Dissociation between early and late components of the event-related brain potential. Brain Research, 1392, 62–79. https://doi.org/10.1016/j.brainres.2011.03.071

Osterhout, L., & Mobley, L. (1995). Event-related brain potentials elicited by failure to agree. Journal of Memory and Language, 34(6), 739–773.

Palolahti, M., Leino, S., Jokela, M., Kopra, K., & Paavilainen, P. (2005). Event-related potentials suggest early interaction between syntax and semantics during on-line sentence comprehension. Neuroscience Letters, 384(3), 222–227. https://doi.org/10.1016/j.neulet.2005.04.076

Philipp, M., Bornkessel-Schlesewsky, I., Bisang, W., & Schlesewsky, M. (2008). The role of animacy in the real time comprehension of Mandarin Chinese: Evidence from auditory event-related brain potentials. Brain and Language, 105(2), 112–133. https://doi.org/10.1016/j.bandl.2007.09.005

Popov, S., & Bastiaanse, R. (2018). Processes underpinning gender and number disagreement in Dutch: An ERP study. Journal of Neurolinguistics, 46, 109–121. https://doi.org/10.1016/j.jneuroling.2018.01.001

Procházka, S., & Gabsi, I. (2017). Agreement with Plural Heads in Tunisian Arabic: The Urban North. In V. Ritt-Benmimoun (Ed.), Tunisian and Libyan Arabic Dialects Common Trends – Recent Developments – Diachronic Aspects (pp. 239–260). Zaragoza: Prensas de la Universidad de Zaragoza.

Ritt-Benmimoun, V. (Ed.). (2017). Tunisian and Libyan Arabic dialects: Common trends, recent developments, diachronic aspects (1a. edición). Zaragoza: Prensas de la Universidad de Zaragoza.

Sasse, H.-J. (1993). Syntactic categories and subcategories. In J. Jacobs, A. von Stechow, W. Sternefeld, & T. Vennemann (Eds.), Syntax. Ein internationales Handbuch zeitgeno ssischer Forschung/An International Handbook of Contemporary Research (Vol. 1, pp. 646–686). Berlin: de Gruyter.

Silverstein, M. (1976). Hierarchy of features and ergativity. In R. M. W. Dixon (Ed.), Grammatical categories in Australian languages (pp. 112–171). New Jersey, NJ: Humanities Press.

Steele, S. (1978). Word order variation: A typological study. In J. H. Greenberg, C. A. Ferguson, & E. A. Moravcsik (Eds.), Universals of human language, vol. 4: Syntax (Vol. 4, pp. 585–623). Stanford University Press: Walter de Gruyter.

Szewczyk, J. M., & Schriefers, H. (2011). Is animacy special? ERP correlates of semantic violations and animacy violations in sentence processing. Brain Research, 1368, 208–221. https://doi.org/10.1016/j.brainres.2010.10.070

Szewczyk, J. M., & Schriefers, H. (2013). Prediction in language comprehension beyond specific words: An ERP study on sentence comprehension in Polish. Journal of Memory and Language, 68(4), 297–314. https://doi.org/10.1016/j.jml.2012.12.002

Tanner, D., & van Hell, J. G. (2014). ERPs reveal individual differences in morphosyntactic processing. Neuropsychologia, 56, 289–301. https://doi.org/10.1016/j.neuropsychologia.2014.02.002

van de Meerendonk, N., Kolk, H. H. J., Chwilla, D. J., & Vissers, C. Th. W. M. (2009). Monitoring in Language Perception. Language and Linguistics Compass, 3(5), 1211–1224. https://doi.org/10.1111/j.1749-818X.2009.00163.x

White, A., Malt, B. C., & Storms, G. (2017). Convergence in the Bilingual Lexicon: A Pre-registered Replication of Previous Studies. Frontiers in Psychology, 7. https://doi.org/10.3389/fpsyg.2016.02081

Wicha, N. Y. Y., Moreno, E. M., & Kutas, M. (2004). Anticipating Words and Their Gender: An Event-related Brain Potential Study of Semantic Integration, Gender Expectancy, and Gender Agreement in Spanish Sentence Reading. Journal of Cognitive Neuroscience, 16(7), 1272–1288. https://doi.org/10.1162/0898929041920487

Wunderlich, D. (2015). Grammatical Agreement. In J. D. Wright (Ed.), International Encyclopedia of Social and Behavioral Sciences (Second Edition, pp. 316–323). Elsevier.

Zawiszewski, A., & Friederici, A. D. (2009). Processing canonical and non-canonical sentences in Basque: The case of object–verb agreement as revealed by event-related brain potentials. Brain Research, 1284, 161–179. https://doi.org/10.1016/j.brainres.2009.05.099

Zawiszewski, A., Santesteban, M., & Laka, I. (2016). Phi-features reloaded: An event-related potential study on person and number agreement processing. Applied Psycholinguistics, 37(3), 601–626.

